# simSOMA: a cell-lineage based simulator of the somatic VAF spectrum in plants

**DOI:** 10.64898/2026.06.28.735079

**Authors:** Frank Johannes

**Affiliations:** Technical University of Munich, Plant Epigenomics, Emil-Ramann-Str. 4, 85354 Freising, Germany; Centre for Research in Agricultural Genomics (CRAG), CSIC-IRTA-UAB-UB, Campus UAB, Bellaterra, Barcelona, Spain

**Keywords:** somatic mutation, somatic epimutation, variant allele frequency, shoot apical meristem, cell lineage, plant development, branching, simulation, developmental bottleneck, clonal mosaicism, plants

## Abstract

Plants accumulate somatic mutations during growth, some of which spread into branches, organs, or reproductive tissues. These variants are of growing interest because they underlie bud-sport traits in perennial crops, contribute to within-organism selection, and are a source of heritable genetic variation. Recent genomic sequencing of bulk or layer-enriched plant tissues has revealed complex variant allele-frequency (VAF) spectra. Interpreting these spectra requires modelling how somatic mutations are filtered or amplified by shoot growth, branching, and organ formation, a problem that is difficult to derive analytically. Here, we present simSOMA, a modular simulator linking plant topologies to shoot apical meristem cell-lineage dynamics. simSOMA models mutation accumulation during stem-cell self-renewal, clonal expansion to the meristem periphery, branch founding, and organ formation. We demonstrate how simSOMA isolates developmental processes that shape organ-level VAF spectra and among-organ variant sharing, and transforms spectra for bulk or layer-enriched sampling, phased or unphased readouts, and read-level sequencing noise. SAM layers can be simulated as independent cell lineages and combined into bulk tissue samples. simSOMA can be used to explore developmental hypotheses about the origin of somatic variation, and to design somatic sequencing experiments and variant-detection strategies. simSOMA is available at https://github.com/jlab-code/simSOMA.

---

Somatic mutations are an unavoidable consequence of plant growth. Once they arise, their fate depends on the cell lineages in which they occur. Many de novo mutations remain confined to small sectors, whereas others can be carried into branches, organs, tissues used for clonal propagation, or reproductive tissues. Although somatic variation in plants has long been studied, modern sequencing approaches, improved genome assemblies, and renewed theoretical interest in somatic evolution have made it possible to revisit these variants at genomic scale [Schoen and Schultz, 2019]. This renewed interest reflects the diverse consequences of somatic variants: they can underlie bud-sport traits in crops, generate somatic mosaicism within long-lived individuals, contribute to within-organism adaptive processes, and provide genetic variation that may enter clonal or sexual lineages [Foster and Aranzana, 2018, Reusch et al., 2021, Schoen and Schultz, 2019].

Much of the above-ground plant body is generated post-embryonically. New stems, leaves, flowers, and lateral branches are produced iteratively from shoot apical meristems (SAMs), allowing plant architecture to expand throughout life. This mode of development is especially pronounced in long-lived perennials such as trees, where repeated branching over years to centuries produces large, hierarchically organized shoot systems. As a consequence, the developmental history of a plant is recorded not only in its physical branching architecture, but also in the cell lineages that have propagated through this architecture [Johannes, 2025b, Tomimoto et al., 2025].

The SAM contains a small population of apical stem cells (ASCs) located near the meristem center [Lyndon, 1998, Kwiatkowska, 2008, Burian et al., 2016]. In the classical anatomical literature these cells are referred to as apical initials or initial cells, and in molecular studies as the stem cells of the central zone; here we use ASC throughout. These cells self-renew (i.e. self-maintain) while producing descendants that are displaced toward the SAM periphery [Burian et al., 2016]. During self-renewal, the ASC population persists continuously through cell division, without being temporarily lost and re-established. Peripheral descendants contribute to organ formation and, during axillary meristem initiation, to the founding of new lateral branches. There are two views of axillary meristem initiation. In the detached-meristem model, a small group of relatively quiescent meristematic cells becomes separated from the surrounding, actively dividing cells as a leaf primordium develops, and is later incorporated into the axillary meristem; in the de novo induction model, peripheral cells proliferate without an early distinction between future axillary-meristem cells and their neighbours, and a subset of them later acquires meristematic identity [Burian et al., 2016, Shi et al., 2016, Wang and Jiao, 2018]. From a cell-phylogenetic perspective, both views are equivalent. In both, only a small number of precursor cells, recruited from a localized region of the SAM periphery, establishes the new lateral meristem. Branching therefore imposes a developmental bottleneck on the cell lineages transmitted from one meristem to the next.

The cellular organization of the SAM adds another level of structure. In seed plants, the SAM is commonly described by a tunica–corpus organization. The outer tunica layers, often referred to as L1 and L2, divide predominantly anticlinally, whereas the inner corpus, or L3, divides in multiple orientations [Esau, 1965, Lyndon, 1990]. These layers can maintain partly independent cell-lineage histories and contribute differently to mature tissues. The L1 typically gives rise to the epidermis, L2 to subepidermal tissues and, in many angiosperms, reproductive lineages, and L3 to internal tissues including vasculature. This layer structure is not universal across all plants, and comparative reviews of shoot-apex organization have long shown that SAM structure varies among major plant groups and taxa [Gifford and Corson, 1971, Gola and Banasiak, 2016]. In particular, the shoot apices of most gymnosperms lack a stable tunica–corpus organization, and cell lineages are replaced between apical zones more frequently than in angiosperms [Esau, 1965, Gifford and Corson, 1971]. Where such layers are developmentally stable, they provide a useful framework for interpreting the cell-lineage fate of somatic variants.

Somatic mutations are observed only through the tissues descended from the layer and lineage in which they arose. Mutations arising in differentiating non-meristematic cells are generally restricted to small local sectors. By contrast, mutations arising in ASCs or their long-lived meristematic descendants can be propagated through branch and organ formation, sometimes becoming fixed within a SAM layer, an organ, a branch, or a larger sector of the plant body [Foster and Aranzana, 2018]. However, fixation is not all-or-none at the level of sampled tissue. The frequency at which a mutation is observed in a sequenced tissue sample depends on when and where it arose, whether its lineage persisted in the ASC niche, which founder lineages were sampled during branch and organ formation, how much subsequent organ growth occurred, and which SAM layers contributed to the sequenced tissue [Johannes, 2025b].

Recent sequencing of bulk and layer-enriched plant tissues has shown that de novo somatic variants can span complex variant allele frequency (VAF) spectra [Xie et al., 2016, Schmid-Siegert et al., 2017, Plomion et al., 2018, Hanlon et al., 2019, Wang et al., 2019, Hofmeister et al., 2020, Orr et al., 2020, Duan et al., 2022, Perez-Roman et al., 2022, Satake et al., 2023, Schmitt et al., 2022, Goel et al., 2024, Amundson et al., 2025, Davis et al., 2026, Meyer et al., 2025, Ren et al., 2021, Yu et al., 2024]. For a somatic variant, the VAF is the fraction of sequencing reads, or, in an idealized cell sample, the fraction of sampled alleles or cells that carry the variant. The VAF spectrum is the distribution of these frequencies across somatic variants in a sample. A sampled organ may contain high-frequency variants inherited from earlier stem-cell or branch lineages, intermediate-frequency variants produced by partial founder contributions or incomplete lineage fixation, and low-frequency private variants generated during terminal organ development. Layer-enriched sequencing can help assign such spectra to particular SAM layers by targeting tissues with known developmental origins [Goel et al., 2024, Amundson et al., 2025]. In contrast, bulk sequencing of whole organs produces a composite spectrum in which layer-specific VAFs are mixed according to the relative contributions of L1-, L2-, and L3-derived cells [Davis et al., 2026].

The organ-level VAF spectrum is therefore an integrated readout of linked plant architecture and cell-lineage dynamics. It reflects mutation accumulation during cell division, ASC self-renewal and turnover, clonal expansion from the ASC niche to the SAM periphery, precursor recruitment during branch and organ formation, terminal organ growth, layer composition, sequencing depth, and the topology connecting sampled organs. Interpreting these spectra therefore requires a developmental model that connects cellular lineage dynamics to the physical plant architecture [Johannes, 2025b,a, Tomimoto et al., 2025].

These developmental constraints are not restricted to de novo genetic variation but also apply to somatic epigenetic variation [Chen et al., 2024]. For instance, stochastic gains and losses of DNA cytosine methylation can arise during mitotic cell divisions and, once established, can be inherited through subsequent cell divisions [Johannes and Schmitz, 2019]. These so-called spontaneous epimutations occur at substantially higher rates than DNA sequence mutations and can generate abundant cell-to-cell epigenetic heterogeneity within the SAM and its descendant tissues [Chen et al., 2024, Vo et al., 2026]. Because genetic mutations and epimutations are transmitted through the same cell lineages, their spatial distribution is shaped by the same developmental processes [Johannes, 2024, Zhou et al., 2025]. A cell-lineage framework for somatic variation is therefore relevant to both plant genomics and plant epigenomics.

A number of theoretical approaches have sought to connect plant development to the fate of somatic (epi)mutations and, more recently, to the VAF spectra observed in sequencing data. Previous studies have modeled cell-lineage behavior in the SAM, including lineage displacement, drift, and selection, as well as cell-lineage sampling during branching [Klekowski and Kazarinova-Fukshansky, 1984a,b, Klekowski et al., 1985, Iwasa et al., 2023, Tomimoto and Satake, 2023, Tomimoto et al., 2025, Chen et al., 2024, Otto and Orive, 1995, Pineda-Krch and Fagerstrom, 1999, Folse and Roughgarden, 2012]. Recent work has also examined how cell-lineage dynamics interact with plant branching architecture to shape mutational diversity within the plant body [Johannes, 2025b,a, Tomimoto et al., 2025]. Some frameworks embed such models within inference procedures to estimate mutation rates or developmental parameters from sequencing data [Johannes, 2025b, Tomimoto and Satake, 2026, Grecu et al., 2025, Tomimoto and Satake, 2023, Shahryary et al., 2020]. However, most existing models retain simplified developmental assumptions, partly because the underlying processes are analytically challenging to describe. As a result, they often use idealized branching structures or focus on selected components of the overall developmental process. It therefore remains difficult to ask how alternative developmental assumptions jointly shape organ-level VAF spectra and among-organ variant sharing across diverse plant topologies.

Whole-plant models illustrate these simplifications. Tomimoto and Satake [2023, 2026] adopted the stochastic-meristem model, in which all *m* initials are replaced after every division cycle by a random sample of their descendants [Klekowski and Kazarinova-Fukshansky, 1984b]. For the small niches typical of seed plants, this rule removes at least one parental lineage in most cycles (with probability 0.87 for *m* = 5), the amount of turnover is fixed by *m* rather than set by a separate parameter, and random resampling ignores the spatial constraints on cells at the SAM

center [Burian and Johannes, 2026]. Yu et al. [2024] treated symmetric (Moran-type) and asymmetric stem-cell maintenance as alternative regimes rather than as division outcomes that occur with graded frequency in the same niche. Our own earlier model allowed stem-cell lineage turnover only at branch-founding events and described the strength of each founding bottleneck phenomenologically, rather than deriving it from the cellular steps of branch initiation [Johannes, 2025a]. None of these models represents the steps between the niche and a sequenced organ explicitly: clonal propagation to the SAM periphery, recruitment of founder cells, organ growth and the sequencing assay.

To address these limitations, we introduce simSOMA, a modular simulator for somatic VAF spectra in plants. simSOMA implements each developmental step as a separate module, and the strength of a branching bottleneck emerges from quantities that can in principle be measured by histology or live imaging: the niche size, the size of the amplified peripheral population and the number of recruited precursors. simSOMA can operate on any user-defined plant topology that can be represented as a rooted acyclic graph, allowing arbitrary branching structures, branch lengths, and sampled organ positions to be linked to explicit cell-lineage dynamics. Topologies can be written by hand, converted from simple branch and organ tables, or derived from terrestrial laser scans of whole tree crowns (Supplementary Text, Section 2). The simulator generates organ-level mutation-count and VAF outputs from developmental processes including ASC self-renewal, clonal expansion from the ASC niche to the SAM periphery, branch founding, organ formation, and sequencing subsampling. These developmental VAF spectra can then be transformed post hoc into observation-specific VAF spectra that reflect layer-specific sampling, bulk tissue composition, phased or unphased readout, and read-level sequencing noise. Here, phased readout refers to variant calls made against a haplotype-resolved (phased) genome assembly, in which a heterozygous somatic variant fixed in all sampled cells is observed at VAF one on its haplotype; unphased readout refers to calls against a collapsed (haploid-merged) diploid assembly, in which reads from both homologous chromosomes are pooled and the same variant is observed at VAF 1*/*2. This separates the developmental processes that generate somatic mosaicism from the observational assumptions that determine how this mosaicism appears in sequencing data.

The simulator is organized into developmental modules (Fig. 1). This modular design makes it possible to vary individual developmental assumptions while keeping the overall traversal of the plant topology fixed. Using simSOMA, we show how individual developmental processes can be isolated, varied, and combined to assess their effects on organ-level VAF spectra and among-organ variant sharing.

**Figure 1:**
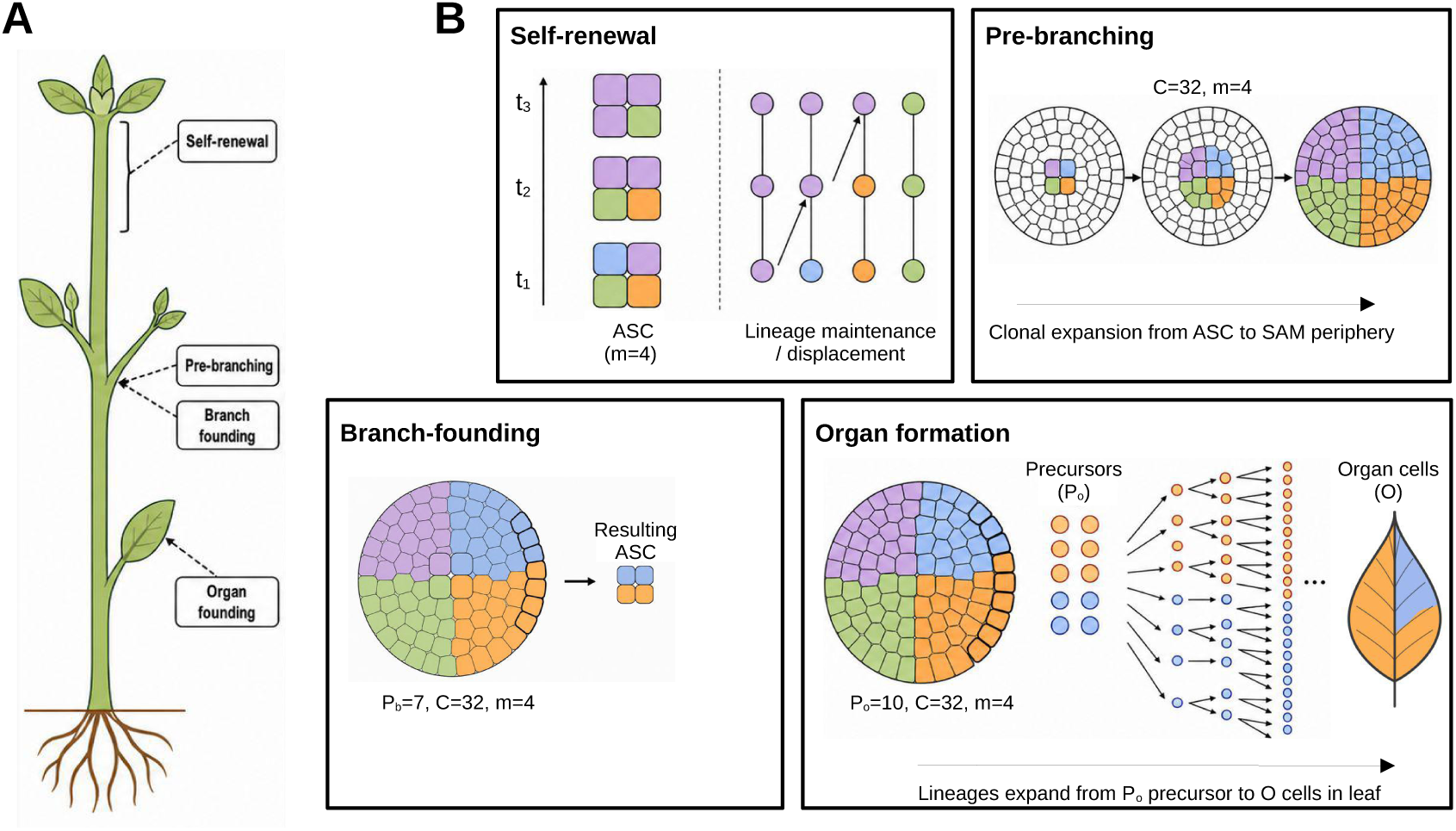
Overview of simSOMA developmental modules. **(A)** Schematic plant topology showing where the developmental modules are active. Self-renewal acts along branch-axis segments between developmental events. Pre-branching clonal amplification and branch founding occur at branch initiation sites, where a lateral branch is produced from the parental shoot apical meristem (SAM). Organ formation occurs at terminal organ-founding sites. **(B)** Schematic representation of the four generative modules. In the self-renewal module, an apical stem-cell (ASC) niche *m* is maintained through time while lineage maintenance and displacement determine which ASC lineages persist in long-lived niche positions. In the pre-branching module, the current ASC state is clonally expanded from the niche toward the SAM periphery, producing a sectorized peripheral cell population *C*. In the branch-founding module, a contiguous peripheral precursor block *P_b_* is recruited from this sectorized SAM population and used to seed the ASC niche of a new lateral branch. In the organ-formation module, a contiguous set of organ precursors *P_o_* is recruited from the SAM periphery and expanded by lineage growth to form a terminal organ consisting of *O* total cells.

## Simulator framework

simSOMA links a user-defined rooted plant topology to explicit cell-lineage dynamics in shoot apical meristems. The topology specifies branch relationships, branch lengths, and the positions of branch and organ events, while the developmental model determines how somatic mutations arise during mitotic divisions and how mutant lineages are transmitted through branching and organ formation. The goal is to generate organ-level mutation counts and VAF spectra, together with summaries of whether variants are private to individual organs (observed in exactly one sampled organ) or shared among organs (observed in two or more). Full mathematical and implementation details are given in the Supplementary Text.

The simulator is organized around four generative modules that correspond to the developmental processes shown in Fig. 1. First, a self-renewal module advances an ordered apical stem-cell niche of size *m* along each branch axis. During self-renewal, cells divide, new mutations are introduced according to a per-division mutation rate, and stochastic lineage displacement can replace one niche lineage by another. The turnover parameter *ρ* controls the frequency of such displacement events and therefore the balance between long-term lineage persistence and lineage turnover within the niche [Burian and Johannes, 2026]. Second, before branch or organ founding, the current niche state is clonally expanded from the stem-cell positions toward the SAM periphery, producing a sectorized peripheral population of size *C*. Third, the branch-founding module recruits a contiguous block of *P_b_* precursor cells from this peripheral population to seed the apical stem-cell niche of a new lateral branch. Fourth, the organ-formation module recruits *P_o_* organ precursors and expands them to form a terminal organ of size *O*. For branch and organ events, the focal sampling position at the SAM periphery can be chosen uniformly or assigned by optional phyllotactic annotation, which maps branch-local developmental sites to angular positions around the meristem. These modules provide the developmental link between mutations carried by meristematic lineages and the VAF spectra observed in sampled organs. Throughout, a developmental bottleneck denotes any step at which a new cell population (a lateral-branch niche or an organ) is founded by a small number of recruited cells, and the developmental depth of a sampled organ denotes the number of branch-founding events on the path from the root of the topology to that organ.

A central feature of simSOMA is that branch-axis progression and local developmental expansion are kept distinct. Branch lengths, given in user-defined topology units such as years or meters, are converted to expected numbers of self-renewal rounds. The corresponding per-division mutation rate is scaled so that the expected mutation input per topology unit remains fixed. In contrast, local amplification during branch founding and organ growth adds mitotic divisions without advancing branch-axis time. This distinction makes it possible to vary stem-cell turnover, branch and organ bottlenecks, terminal organ size, and topology independently while preserving a consistent interpretation of developmental time. Details of the topology mapping, phyllotactic event annotation, module-specific state transitions, and time semantics are provided in Supplementary Text Sections 2–8.

Mutations arise at a per-division rate *µ*_div_ in all modules. Optionally, the rate during organ amplification can be multiplied by a factor *g* (default *g* = 1) to represent elevated mutation rates in terminal, differentiating cells [Meyer et al., 2025].

During a simulation, simSOMA traverses the topology branch by branch. Self-renewal propagates branch-local stem-cell state between developmental events, whereas branch-founding and organ-formation events trigger local sampling from the current SAM-periphery state. Optional phyllotactic annotation can assign focal positions to events along a branch, allowing repeated organ or branch initiation to sample from different sectors of the SAM periphery. The simulator exports organ-level mutation counts, lineage-composition information, event metadata, VAF spectra, and private-versus-shared variant summaries. The output tables and downstream VAF transformations, including bulk or layer-enriched sampling and phased or unphased readout, are described in Supplementary Text Sections 8–10. User-settable cross-module and module-specific parameters are summarized in Appendix Tables 1 and 2. simSOMA is implemented in Python (version *≥*3.10; dependencies NumPy, pandas and Matplotlib), is installable with pip, and is run from configuration files through a command-line interface (e.g. simsoma run config.json). The simulator and a tutorial for running it can be found at https://github.com/jlab-code/simSOMA and https://github.com/jlab-code/simSOMA-tutorial, respectively.

## Results

We applied simSOMA across a series of growth scenarios to isolate, vary, and combine individual developmental processes. In what follows we highlight several key results.

### Global parameter scan identifies key VAF determinants

In an initial simulation, we perform a global parameter scan across a realistic branching topology, asking how stem-cell dynamics, developmental bottlenecks, and topology jointly shape somatic VAF spectra. The simulated topology contained 20 sampled organs on branches of orders 1 to 5, at different root-to-organ developmental depths (Fig. 2A, Supplementary file 1). Across this topology, we varied parameters controlling stem-cell self-renewal, including the number of stem-cell positions, *m*, and the turnover parameter, *ρ*, as well as parameters controlling developmental bottlenecks during branching and organ formation, including the number of branch precursors, *P_b_*, and organ precursors, *P_o_* (Supplementary file 2).

**Figure 2:**
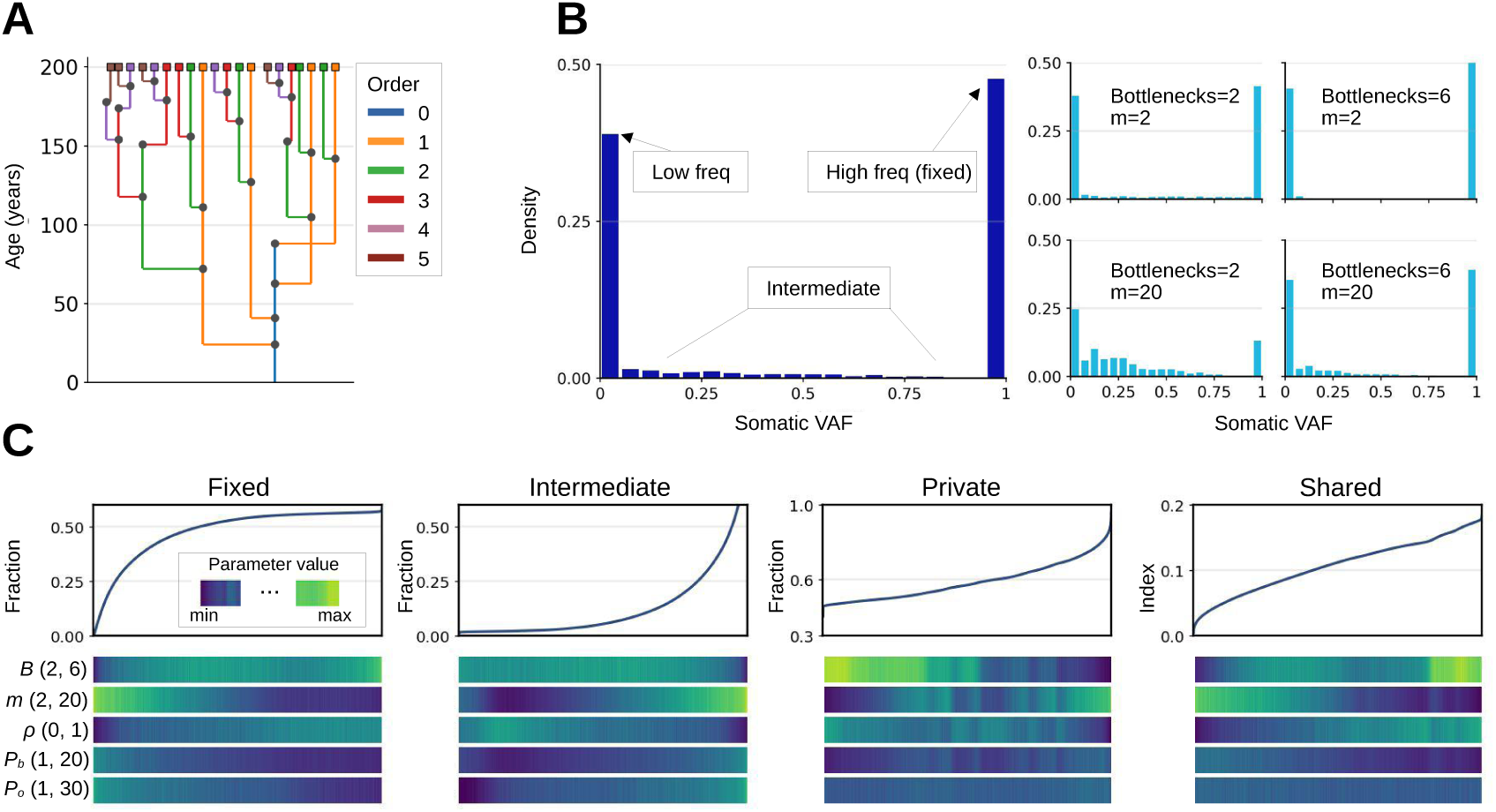
Global parameter scan of developmental effects on somatic VAF spectra. (A) Simulated 200-year branching topology with sampled organs distributed across branch orders 1 to 5. Branch colors indicate branch order. (B) Pooled organ-level VAF spectrum and representative conditional VAF spectra. The pooled spectrum illustrates the private, intermediate, and fixed parts of the VAF distribution, whereas the smaller panels show representative parameter combinations spanning stem-cell niche size and developmental bottleneck settings. “Bottlenecks” denotes the developmental depth of the sampled organ, i.e. the number of branch-founding events on its root-to-organ path (2 = shallow organ with few nested bottlenecks; 6 = deep organ with many nested bottlenecks); it is a property of the organ’s position in the topology, not a simulation parameter, and is labelled *B* in panel C. (C) Ranked global scan of fixed, intermediate, private, and shared statistics. For each statistic, simulations are ordered by the statistic value; the heat strips below each curve show the corresponding relative parameter values for developmental depth (*B*), stem-cell niche size *m*, turnover *ρ*, branch precursor number *P_b_*, and organ precursor number *P_o_*.

The scan was a full factorial grid over *m ∈ {*2, 3, 4, 5, 6, 8, 10, 12, 16, 20*}*, 16 values of *ρ* between 0 and 1, 12 values of *P_b_* between 1 and 20, and 12 values of *P_o_*between 1 and 30, with 50 replicate simulations per parameter combination. The remaining parameters were fixed: *C* = 64, organs of *O* = 10^4^ cells of which 1% (100 cells) were sampled, a mutation rate of one mutation per lineage per year, and five self-renewal rounds per year, with branch lengths converted deterministically into self-renewal rounds. Because only min(*P_b_, m*) recruited cells can seed a new niche (see below), combinations with *P_b_ > m* are developmentally equivalent to *P_b_* = *m*. Configuration files for the simulations of Figs. 2–5 are provided as Supplementary file 2.

**Figure 3:**
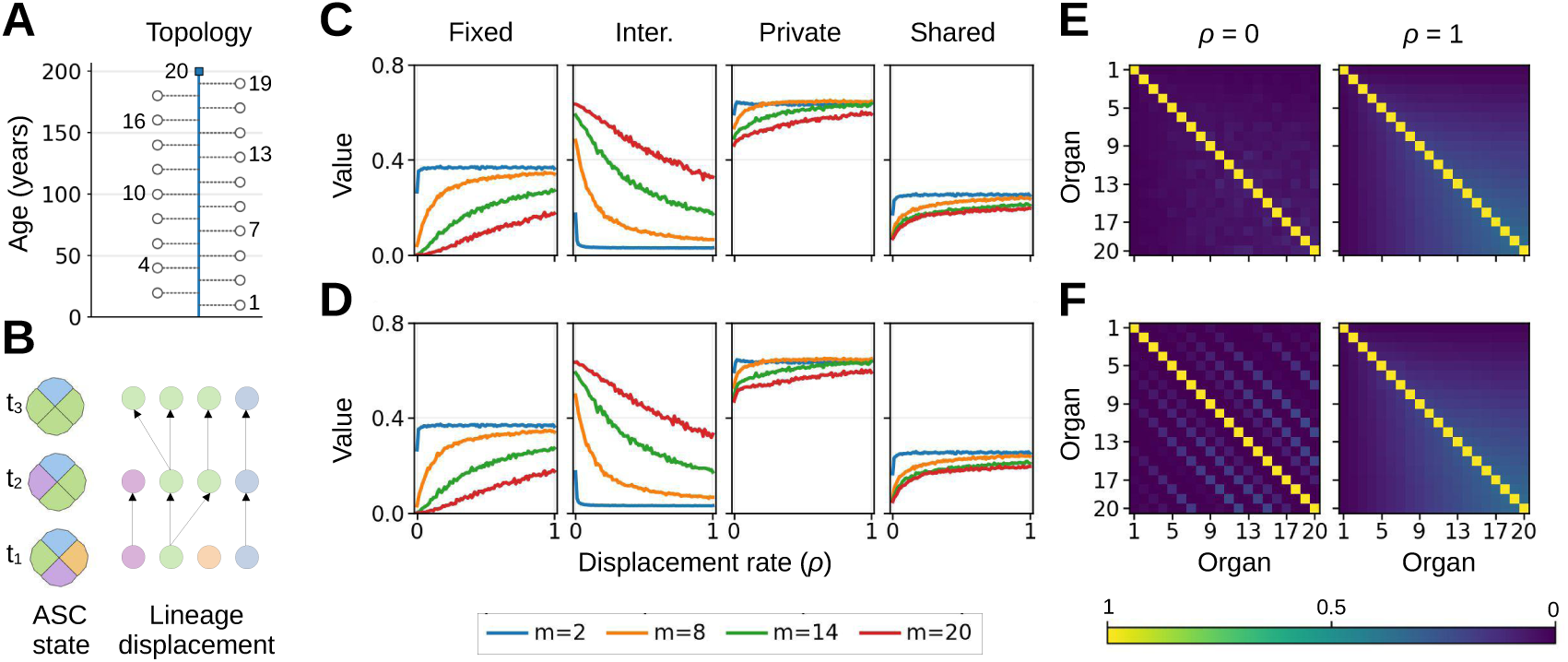
Stem-cell turnover and phyllotaxy shape VAF spectra and organ-level mutation sharing. (A) Single-axis topology with 20 organs sampled at regular positions along a 200-year developmental axis. (B) Conceptual schematic of ASC state and lineage displacement through time. Lineage displacement changes which stem-cell lineages occupy long-lived niche positions and therefore controls the persistence of mutant lineages. (C,D) Marginal VAF and sharing summary statistics as a function of displacement rate *ρ*, without phyllotaxy (C) and with spiral phyllotaxy (D). Panels show fixed fraction, intermediate fraction, private fraction, and normalized sharedness; colors indicate stem-cell niche size *m*. (E,F) Pairwise organ-sharing matrices for *ρ* = 0 and *ρ* = 1, without phyllotaxy (E) and with spiral phyllotaxy (F). Heatmaps show pairwise Jaccard sharing between organs on a common 0–1 color scale.

**Figure 4:**
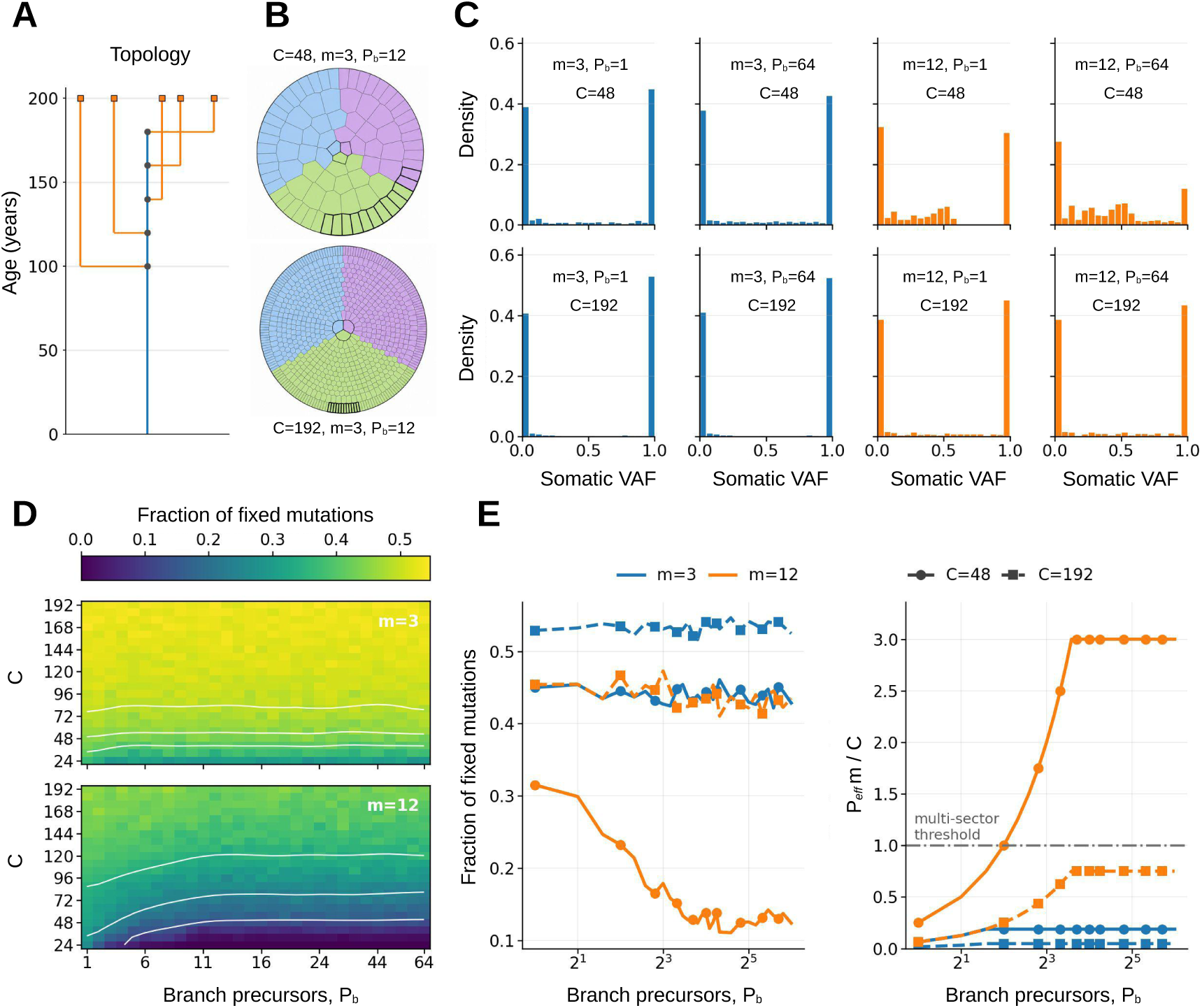
Branch founding depends on the realized precursor footprint relative to peripheral clonal sector size. (A) Simplified branching topology with five lateral branches and one sampled terminal organ per lateral branch. (B) Peripheral SAM sector schematics for two values of pre-branching amplification *C*, with *m* = 3 and requested branch precursor number *P_b_* = 12. Highlighted cells indicate the recruited contiguous branch precursor block. (C) Representative VAF spectra for combinations of stem-cell niche size *m*, requested branch precursor number *P_b_*, and peripheral cell number *C*. (D) Full scan of the fixed-marker fraction across requested branch precursor number *P_b_*and peripheral cell number *C*, shown separately for *m* = 3 and *m* = 12. White contours summarize the broad trend across the heatmap. (E) Left: fixed-marker fraction as a function of requested branch precursor number *P_b_*, showing the plateau that occurs when the realized ASC-founder number reaches the niche-size cap. Right: realized effective branch-founder footprint, *P*_eff_ *m/C*, as a function of requested *P_b_*. The horizontal line marks *P*_eff_ *m/C* = 1, the approximate transition into the multi-sector sampling regime.

**Figure 5:**
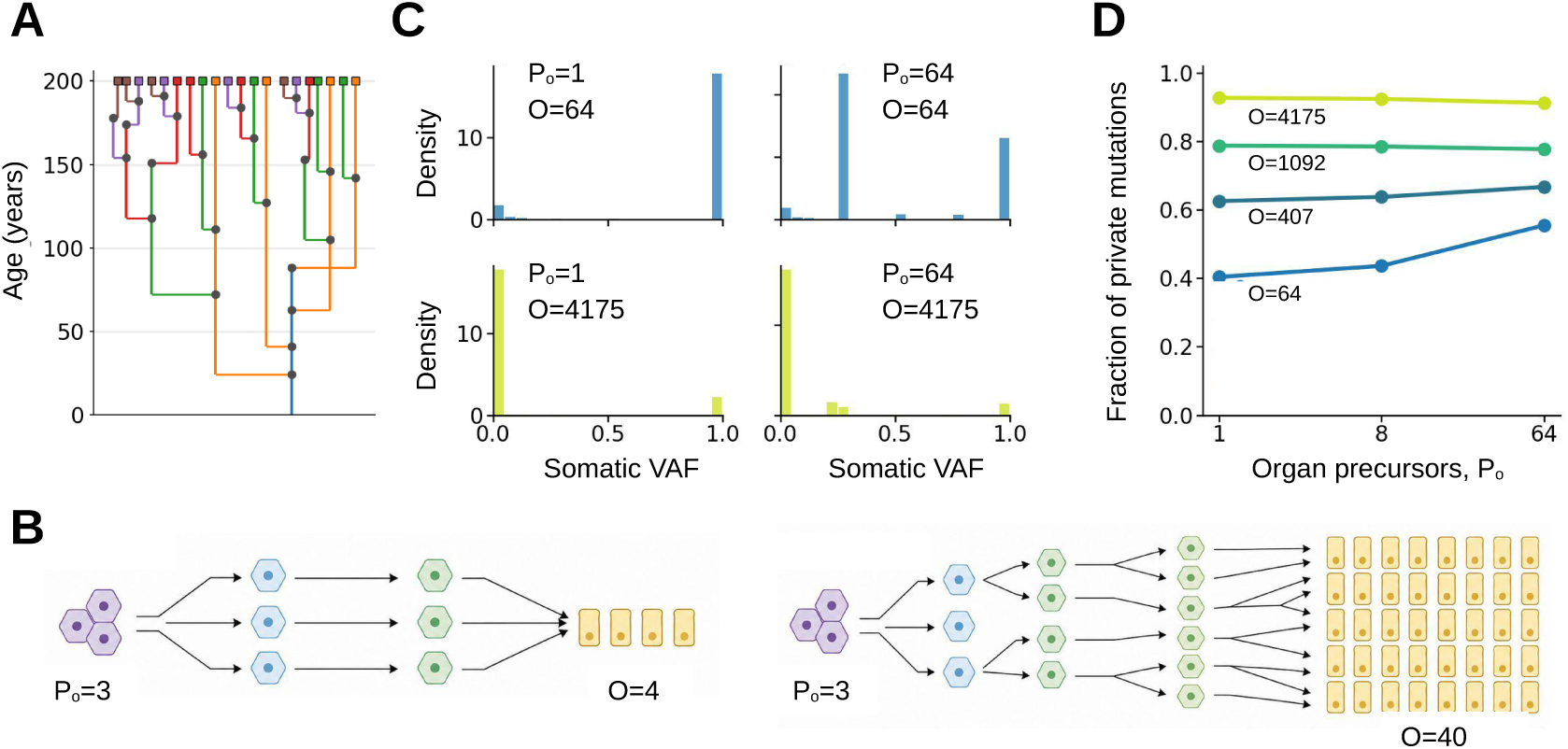
Terminal organ growth generates private low-frequency variants. **(A)** The 200-year multibranch topology used for the organ-formation simulations, with 20 sampled terminal organs distributed across branch orders. **(B)** Conceptual schematic of organ formation. The same number of recruited organ precursors *P_o_* can expand to a shallow or deep terminal organ, represented by different final organ cell numbers *O*. **(C)** Representative somatic VAF spectra for combinations of organ precursor number *P_o_* and final organ cell number *O*. Increasing organ depth produces a large low-VAF component, whereas *P_o_* modulates the severity of the organ-founding bottleneck and the intermediate-frequency component. **(D)** Fraction of private mutations as a function of recruited organ precursor number *P_o_*, stratified by final organ cell number *O*. The private fraction increases strongly with organ cell number and is less sensitive to precursor number once organ-specific expansion is deep.

To compare simulations, we defined four organ-level summary statistics. For a sampled organ *o*, let *M_o_* denote the set of mutations observed in that organ, and let *v_io_* be the sampled VAF of mutation *i* in organ *o*. The fixed fraction was defined as

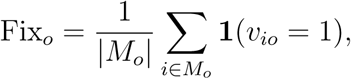

that is, the fraction of observed mutations that are fixed in the sampled cells of organ *o*. The intermediate fraction was defined as

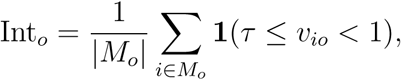

where *τ* = 0.05 was used as the lower VAF threshold. Thus, intermediate variants are non-fixed variants that reach at least 5% VAF in a sampled organ.

We also defined two statistics describing how mutations are distributed across sampled organs. Let *D_i_* be the sharing degree of mutation *i*, defined as the number of sampled organs in which that mutation is observed. The private fraction of organ *o* was defined as

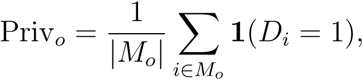

that is, the fraction of mutations observed in organ *o* that are unique to that organ. Finally, sharedness was defined as the normalized mean sharing degree of mutations observed in organ *o*:

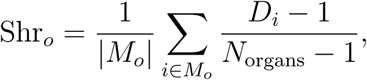

where *N*_organs_ is the number of sampled organs. This statistic equals 0 when all variants in an organ are private and equals 1 when all variants observed in that organ are present in every sampled organ. Importantly, these statistics describe two different aspects of the simulation output. Fixed and intermediate fractions describe the within-organ VAF state of mutations, whereas private fraction and sharedness describe the distribution of mutation across organs. These classes are therefore not mutually exclusive. For example, a mutation can be fixed within one organ and also shared across several organs.

The pooled VAF spectrum was strongly enriched near the extremes of the VAF distribution, with many low-frequency variants and a substantial class of variants fixed within sampled organs (Fig. 2B). This pooled pattern, however, masked substantial parameter sensitivity. Conditional VAF spectra stratified by ASC niche size *m* and the number of branching bottlenecks along root-to-organ paths showed that organs behind many nested bottlenecks (developmental depth 6) combined with small *m* produced a more pronounced bimodal spectrum, with increased density near low VAFs and fixation. By contrast, organs behind few bottlenecks (depth 2) combined with larger *m* maintained more variants at intermediate frequencies.

The ranked global scan summarized these effects across all parameter combinations (Fig. 2C). Fixed and intermediate fractions varied in opposite directions: parameter combinations that increased lineage sampling and developmental drift, including low *m*, high *ρ*, low *P_b_*, low *P_o_*, or greater developmental depth, increased fixation, whereas larger stem-cell and precursor pools maintained more variants at intermediate VAFs. In addition to these within-organ VAF effects, the scan revealed strong effects on how mutations were distributed across organs. The private and shared statistics depend on the topology through the nesting of bottlenecks. Each branch-founding event partitions the plant into sub-trees, and a mutation can be shared only by organs that descend from the niche in which it arose. Deeply nested organs therefore accumulate more branch-specific variants, which are private when a branch carries a single sampled organ, whereas variants arising early in the main axis can be shared widely only if they survive turnover and every intervening bottleneck. The effect of branching architecture on private and shared variants is thus captured by the developmental depth of the sampled organs and by the arrangement of sampled organs on branches, as analysed for fixed somatic mutations in Johannes [2025b].

Together, this global parameter scan highlights that somatic VAF spectra are shaped by the combined effects of stem-cell dynamics, developmental bottlenecks, and topology. Because these parameters have overlapping effects in the global scan, we next examined individual components of the model separately to isolate how self-renewal, branching, and organ formation contribute to fixed, intermediate, private, and shared somatic variant classes.

### Stem-cell turnover shapes VAF spectra and phyllotactic sharing

The global parameter scan identified stem-cell self-renewal as a major determinant of somatic VAF patterns. We therefore isolated the effects of stem-cell niche size and lineage turnover in a simplified single-axis topology. In this topology, organs were sampled at regular intervals along a 200-year developmental axis (Fig. 3A, Supplementary file 1). We varied the number of stem-cell positions, *m*, and the lineage-turnover parameter, *ρ* (Supplementary file 2). Again, the parameter *ρ* controls the probability that one stem-cell lineage displaces another during self-renewal, as illustrated by the ASC-state and lineage-displacement schematic (Fig. 3B). We did not examine the impact of displacement bias, as recent theoretical work showed that its effect is marginally small relative to the impact of *ρ* [Burian and Johannes, 2026]. To test whether spatially structured organ recruitment alters somatic mutation patterns, we compared otherwise identical simulations without phyllotaxy and with spiral phyllotaxy (see Supplementary text for a mathematical definition).

We found that the marginal VAF summary statistics were governed primarily by *m* and *ρ*, and were only weakly affected by phyllotaxy (Fig. 3C,D). Smaller stem-cell populations produced higher fixed fractions, consistent with stronger developmental drift in the stem-cell pool. Increasing *ρ* further increased fixation for most values of *m*, while reducing the fraction of intermediate-frequency variants. Thus, stem-cell turnover shifts somatic variants away from long-lived intermediate-frequency states and toward either loss or fixation. Larger values of *m* buffered this effect and maintained a larger fraction of intermediate-frequency variants, consistent with slower developmental drift in a larger stem-cell population. These results agree with earlier models of stem-cell drift in the SAM, in which small niches and frequent lineage replacement accelerate fixation [Klekowski and Kazarinova-Fukshansky, 1984b, Iwasa et al., 2023], and with the use of this drift as a somatic genetic clock in clonal species [Yu et al., 2024]; simSOMA reproduces this behaviour within an explicit topology and adds its consequences for sharing among organs. Note that *m* counts long-lived ASC positions per SAM layer, i.e. initial cells in classical terminology. Clonal analyses and live imaging indicate that typically only 2–4 ASCs per layer persist at the SAM center, because few cells can simultaneously occupy the central domain of minimal cell displacement; larger numbers are possible in elongated meristems such as the ellipsoidal SAM of *Zostera marina* [Burian et al., 2016, Yu et al., 2024, Burian and Johannes, 2026]. We therefore regard *m ≤* 4 as the biologically typical range; values up to *m* = 20 were included to show the large-niche limit and to cover meristems with extended central domains.

When looking across organs, private variants and normalized sharedness both increased with *ρ*, but for different reasons (Fig. 3C,D). In the single-axis topology, organs are sampled sequentially along the same developmental axis. A mutation carried by a stem-cell lineage can therefore be observed in later organs only if that lineage persists in the niche. Increasing *ρ* accelerates lineage replacement.

Mutations carried by lineages that are displaced are removed from future organs and, if sampled before loss, often remain private or narrowly distributed. By contrast, mutations carried by lineages that survive turnover, or rise to dominance, are propagated through a longer subsequent portion of the axis and are therefore shared among more organs. Thus, turnover simultaneously increases the private component and increases the average sharing degree of the persistent component.

Although phyllotaxy had little effect on the marginal summary statistics, it altered the pairwise structure of mutation sharing among organs (Fig. 3E,F). In the absence of phyllotaxy, pairwise sharing was dominated by the temporal organization of the single-axis topology. With spiral phyllotaxy, additional off-diagonal structure appeared in the sharing matrix when lineage turnover was low, *ρ* = 0, indicating repeated spatial sampling of related stem-cell sectors. This phyllotactic signal was strongly reduced when lineage turnover was high, *ρ* = 1, and the sharing matrices with and without phyllotaxy became more similar.

Together, these simulations show that stem-cell niche size and turnover dominate the marginal VAF spectrum, whereas phyllotaxy mainly affects the spatial structure of variant sharing among organs. Spatial organ recruitment therefore leaves its clearest signal when stem-cell lineages remain spatially structured for long enough to be sampled repeatedly. The size of the phyllotactic signal also depends on the sampling design: it is a property of pairs of organs recruited from the same branch axis, so it is diluted when few organs are sampled per axis or when sampled organs are distributed over many branches, and it is strongest for densely sampled single axes as in Fig. 3.

### Branching bottlenecks depend on precursor number relative to clonal sector size

Beyond ASC self-renewal, the global parameter scan identified branch founding as an important determinant of fixed somatic variants. This effect is expected because branch initiation samples only a subset of peripheral SAM cells and can therefore impose a pronounced cell-lineage bottleneck. To examine this process in more detail, we isolated the effects of pre-branching amplification and branch precursor recruitment in a simplified branching topology (Fig. 4A). This topology consists of a main developmental axis with multiple first-order lateral branches, each terminating in a single sampled organ (Supplementary file 1, Supplementary file 2). Branch founding is modelled as the recruitment of a contiguous block of *P_b_* peripheral cells, within which a contiguous block of at most *m* cells becomes the stem-cell niche of the lateral branch; recruited cells are physically connected, so they cannot be drawn independently from across the periphery. Because each sampled organ is separated from the main axis by one branch-founding event, this design isolates branch-level bottlenecks while minimizing additional topological complexity.

Before branch initiation, the peripheral SAM population was expanded to *C* cells. Because these precursors are recruited as a contiguous block, their effect should depend on the size of the block relative to the spatial extent of clonal sectors in the peripheral SAM. The relevant sector-size scale is the average number of peripheral cells contributed by each stem-cell lineage. For a stem-cell population of size *m*, this scale is approximately *C/m* (the exact condition is derived below). Thus, branch founding should preserve high fixation when the realized founder block remains small relative to the clonal sector size. In contrast, when the realized founder block approaches or exceeds *C/m*, branch founding can enter the multisector sampling regime. In this regime, neighboring precursor cells are more likely to span multiple stem-cell-derived sectors, reducing the probability that the branch is founded by descendants of a single lineage. We call a founding event monoclonal if all recruited founders descend from a single clonal sector of the periphery (one niche position at the time of the event) and polyclonal otherwise.

The sector schematics in Fig. 4B show how the same branch precursor block can remain within one large sector or span several smaller sectors, depending on *C* and *m*. The representative VAF spectra and full parameter scan illustrate the consequences of this sampling geometry (Fig. 4C,D). Conditions with small *m*, large *C*, or small requested branch precursor number produced strong fixed-VAF peaks, consistent with branch founding from a restricted clonal sector. For small *m*, clonal sectors were large relative to many recruited precursor blocks, and the fixed fraction therefore remained high across much of the scan. In contrast, larger *m*, smaller *C*, or larger requested branch precursor numbers reduced the average sector size relative to the founder block, increasing the probability of multi-sector sampling and lowering the fixed fraction. Increasing *C* partially buffered this effect by enlarging peripheral clonal sectors before branch initiation. Intermediate-frequency variants were retained under these less restrictive sampling regimes, although they can also include mutations that arose after branch founding as well as variants already present before the bottleneck.

The plateau in the branch-precursor response (Fig. 4E, left) reflects the difference between the requested precursor number *P_b_*, which describes the size of the contiguous branch-founding field, and the realized founder population of the new lateral SAM. Because the new SAM contains only *m* long-lived stem-cell positions, at most *m* precursor-derived cells can seed its niche; additional precursors may contribute to non-ASC branch tissues but do not establish long-lived lineages. The realized founder number is therefore

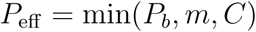

(denoted *P*_b_^eff^ in the Supplementary Text and in Table 2), which equals min(*P_b_, m*) in the parameter range used here (*C ≥ m*). Accordingly, the fixed fraction declines as *P_b_* increases and plateaus once *P_b_* reaches *m*. The right panel of Fig. 4E shows the realized founder footprint relative to the average clonal sector, *P*_eff_ */*(*C/m*) = *P*_eff_ *m/C*. It saturates at *m*^2^*/C*, so only combinations with *m*^2^*/C* above about one enter the multi-sector regime.

This threshold can be made exact. If the focal position of a founding event is placed uniformly on the periphery and *C* is divisible by *m*, a contiguous block of *P*_eff_ founders is polyclonal with probability

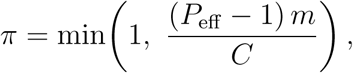

and spans on average 1 + *π* clonal sectors (Supplementary Text, Section 7.5). The term *P*_eff_ *−* 1 rather than *P*_eff_ reflects that a single founder is always monoclonal; for larger *P*_eff_ the two criteria agree. The probability of polyclonal founding therefore depends on precursor number, niche size and peripheral population size only through the compound quantity *ϕ* = (*P*_eff_ *−* 1)*m/C*, and the approximate transition at *P*_eff_ *m/C ≈* 1 in Fig. 4E corresponds to *ϕ ≈* 1. Because *P*_eff_ *≤ m* for branches, *π ≤* (*m −* 1)*m/C*: unless the niche is large relative to the peripheral sectors (*C < m*(*m −* 1)), branch founding is predominantly monoclonal. Organ founding, for which *P_o_* is not capped by *m*, can be strongly polyclonal. Because only *P*_eff_ = min(*P_b_, m*) affects the transmitted lineages, *P_b_* itself cannot be identified from genetic data beyond *P_b_* = *m*.

Together, these simulations show that branch founding is governed by the realized founder footprint relative to peripheral clonal sector size. This explains why fixation declines under multi-sector sampling and why the response plateaus once the lateral SAM founder number is capped by *m*.

### Organ growth is the main source of private variants

The previous simulations revealed that stem-cell dynamics and branch founding shape the VAF spectrum before organ initiation. We next asked how terminal organ formation affects the distribution of somatic variants among sampled organs. To isolate this process, we used the 200-year branched topology with 20 sampled terminal organs distributed across branch orders 1 to 5 (Fig. 5A, Supplementary file 1). Self-renewal and branch-founding parameters were held fixed, with *m* = 4,

*ρ* = 0, *C* = 64, and branch precursor number *P_b_* = 4. We then varied the number of organ precursors, *P_o_*, and the final number of cells in the sampled organ, *O* (Supplementary file 2).

The organ-formation module separates two developmental steps (Fig. 5B). First, a terminal organ is founded from *P_o_* precursor cells. Second, these precursor-derived lineages expand to form the final sampled organ cell population. Thus, *P_o_*controls the severity of the organ-founding bottleneck, whereas *O* controls the depth of subsequent organ-specific cell-lineage expansion. This distinction is important because mutations that occur after organ initiation are restricted to a single terminal organ and therefore contribute directly to the private variant class.

A simple counting argument explains why organ size is expected to dominate the number of private variants. In the current implementation, organ precursors are expanded by balanced binary growth. Each division produces two daughter cells, and each daughter independently acquires new mutations at rate *g µ*_div_. Growing from *P_o_* precursor cells to *O* organ cells requires *O − P_o_* divisions, which produce 2(*O − P_o_*) daughter cells. The expected number of mutations arising during organ growth is therefore

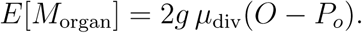

Because these mutations arise after organ initiation, they are restricted to a single organ and are private. The frequency of such a mutation is the fraction of organ cells that descend from the daughter cell in which it arose. That is, a mutation arising in the last round of divisions is carried by a single cell (*q* = 1*/O*), whereas one arising in the first division of a precursor is carried by up to half of that precursor’s descendants (*q ≈* 1*/*(2*P_o_*)). Late divisions vastly outnumber early ones, so private variants are concentrated at low frequency. More precisely, the number of daughter cells whose descendants make up at least a fraction *q* of the organ is about 2(1*/q − P_o_*), so the expected number of private variants with frequency at least *q* is

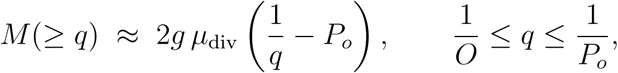

which agrees closely with full simulations (Supplementary Fig. S3). This is the same 1*/q* law that characterizes neutral subclonal mutations in exponentially growing tumours, where the cumulative number of mutations above frequency *q* is linear in 1*/q* with slope proportional to the mutation rate per cell division [Williams et al., 2016, 2018]. Two consequences follow. First, the slope of the low-frequency tail measures the per-division mutation rate during organ growth, 2*gµ*_div_, after rescaling by the source-layer contribution and phasing factor of the observation model (below); comparing it with the mutation rate inferred from shared variants is therefore a direct test for elevated mutation rates in terminal cells [Meyer et al., 2025]. Second, a higher terminal mutation rate (*g >* 1) multiplies the number of private variants at all frequencies in this range, but adds intermediate-frequency variants only through the first few divisions of each precursor (*q* ≲ 1*/*(2*P_o_*)); it therefore mainly deepens the low-frequency tail rather than altering the intermediate part of the spectrum generated by earlier modules.

The representative VAF spectra and full parameter scan show this behavior (see Fig. 5C,D). When the final organ cell number was small, the spectrum was dominated by variants inherited from the organ precursors. Under strong organ-founding bottlenecks, many of these inherited variants reached high or fixed VAFs within the sampled organ, whereas larger *P_o_* mixed more precursor-derived lineages and increased the intermediate-frequency component. Thus, *P_o_* primarily affected the within-organ VAF state of variants already present at organ initiation. In contrast, increasing *O* produced a large excess of low-VAF private variants. The private fraction increased strongly with *O*, and for deep organs remained high across the range of *P_o_*, indicating that most private variants were generated during terminal organ growth rather than inherited from earlier stem-cell or branch lineages.

Together, these simulations show that terminal organ growth is the main source of private, low-frequency variants, whereas organ precursor number mainly modulates the VAF distribution of variants present at organ initiation.

### From developmental to observed VAF spectra

So far we have treated the simulated developmental VAF, *v_io_*, as the directly observed quantity. This corresponds to an idealized observation scenario in which organs are sampled in a layer-specific manner and mutation calls are derived from read alignments to a phased genome assembly. In this case, the sampled cells are treated as descendants of a single SAM layer, and a mutation fixed within that sampled layer has observed VAF one. However, most empirical sequencing designs differ from this idealization. Bulk tissue samples can combine cells derived from multiple histogenic layers, and unphased or collapsed diploid assemblies report heterozygous somatic variants relative to both homologous chromosomes. Under such measurement conditions, a mutation that is fixed within its developmental source layer need not appear at VAF one in the sequencing data (Fig. 6).

**Figure 6:**
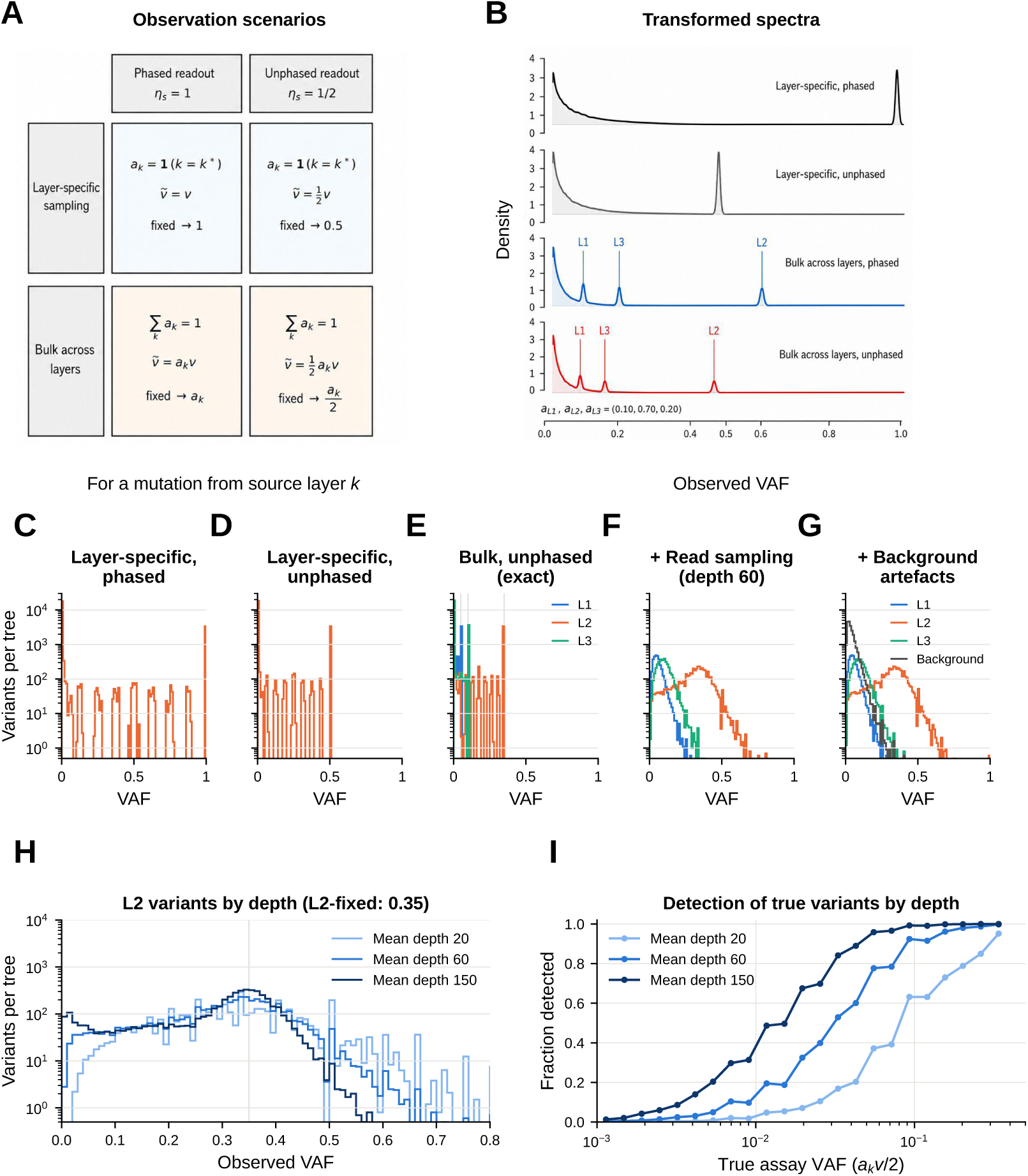
Observation-model transformations of developmental VAF spectra. **(A)** Four observation scenarios: layer-specific or bulk sampling, combined with phased or unphased readout. The observed VAF of a mutation from source layer *k* is its developmental VAF multiplied by the layer contribution *a_k_* (*a_k_* = 1 for the sampled layer under layer-specific sampling; Σ*_k_ a_k_* = 1 in bulk samples) and by the phasing factor *η_s_* (1 phased, 1*/*2 unphased). **(B)** Schematic spectra for the same developmental spectrum under the four scenarios: layer-fixed mutations appear at 1 (layer-specific, phased), 1*/*2 (layer-specific, unphased), *a_k_* (bulk, phased) or *a_k_/*2 (bulk, unphased); example contributions (*a*_L1_*, a*_L2_*, a*_L3_) = (0.10, 0.70, 0.20). **(C–I)** Simulated spectra for ten trees on the topology of Fig. 2A (*m* = 4, *ρ* = 0, *C* = 64, *P_b_* = 4, *P_o_* = 8, *O* = 10^4^ cells with 10^3^ sampled, one mutation per lineage per year in each of three independently simulated layers; variants per tree, organs pooled). **(C)** L2, layer-specific, phased. **(D)** Same, unphased. **(E)** Bulk unphased sample, exact transformation, coloured by source layer; grey lines mark *a_k_/*2. **(F)** After read sampling (calibrated depth model, mean depth 60; variant retained if *≥* 2 variant reads in at least one organ). **(G)** As F, with 2000 background artefact sites. **(H)** Observed VAFs of L2 variants at mean depths 20, 60 and 150. **(I)** Fraction of true variant–organ occurrences detected, as a function of their true (assay) VAF *a_k_v/*2.

We therefore distinguish four simple observation scenarios: layer-specific sampling with phased readout, layer-specific sampling with unphased readout, bulk sampling across layers with phased readout, and bulk sampling across layers with unphased readout (Fig. 6A). These scenarios differ only in how the same underlying developmental VAF is converted into a measured VAF. To make these observational effects explicit, simSOMA can transform exact simulated VAF spectra into user-defined observation scenarios without rerunning the developmental simulation.

Let *v_io_*^(*k*)^ denote the developmental VAF of mutation *i* in organ *o*, conditional on mutation *i* having arisen in source layer *k*. For an observation scenario *s*, let *a_k_*^(*s*)^ denote the contribution of source layer *k* to the sampled tissue. For layer-specific sampling of layer *k^∗^*,

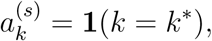

whereas for bulk sampling across layers the layer contributions satisfy

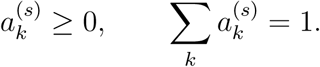

We further define a phasing factor, assuming a diploid species,

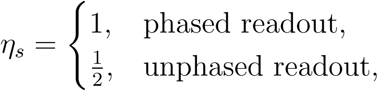

where the unphased factor reflects that a heterozygous somatic variant fixed in all sampled cells is expected at VAF 1*/*2 when reads are counted against both homologous chromosomes. For a mutation arising in source layer *k*, the observed VAF under scenario *s* is therefore

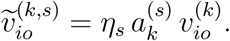

This expression shows that the observation model does not change the underlying developmental frequency *v_io_*^(*k*)^. Rather, it rescales that frequency by the contribution of the source layer to the sampled tissue and by whether the variant is measured in a phased or unphased representation (Fig. 6A).

To obtain the source-layer frequencies *v_io_*^(*k*)^, simSOMA simulates the SAM layers as independent cell-lineage histories on the same plant topology: one simulation per layer, each with its own random seed and, optionally, its own mutation rate, and with shared developmental parameters. The topology, including phyllotactic event positions, is shared, so that branch and organ initiation sites occupy the same angular positions in all layers. Because we assume that there is no cell exchange between layers, every mutation arises in, and can become fixed in, exactly one layer. The bulk VAF of a mutation of layer *k* in organ *o* is then η_s_*a_ko_*^(*s*)^v*_io_*^(*k*)^, where layer contributions can differ among organs. This replaces the post-processing approximation of the preprint, in which one layer-equivalent spectrum was copied into each layer, so that all layers shared an identical lineage history.

These transformations have direct consequences for the interpretation of VAF spectra (Fig. 6B). Under layer-specific phased observation, a layer-fixed mutation appears at

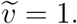

Under layer-specific unphased observation, the same mutation appears at

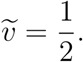

Under bulk phased observation, a mutation fixed in source layer *k* appears at

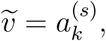

and under bulk unphased observation it appears at

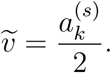

### Thus, for a three-layer bulk sample with contributions

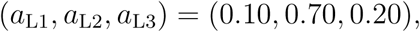

source-layer-fixed mutations are expected at VAFs 0.10, 0.70, and 0.20 under phased bulk observation, and at VAFs 0.05, 0.35, and 0.10 under unphased bulk observation (Fig. 6B). Peaks or enrichments at these values therefore reflect the observational mapping from developmental fixation to measured VAF, not different developmental fixation states.

### Read-level observation

The transformations above give the expected VAF of each mutation in a sequenced sample. Empirical VAFs are additionally affected by finite and variable sequencing depth, read sampling, sequencing and alignment artefacts, and the variant-calling procedure. simSOMA therefore allows users to add a read-level transformation, implemented as a separate, versioned observation model (Supplementary Text, Section 10). For each site *i* and sampled organ *o*, a read depth *D_io_* is drawn and the number of variant reads is *A_io_ ∼* Binomial(*D_io_, p_io_*) with *p_io_* = *ṽ_io_*(1 *−e*) +(1 *− ṽ_io_*)*e*, where *ṽ_io_* is the expected

VAF given by the transformations above and *e* is the sequencing error rate; a beta-binomial can be used instead to represent overdispersion. Depth can be fixed, normally or Poisson distributed, or follow a calibrated model. In the calibrated model, read depth at site *i* in organ *o* is Poisson with mean *D u_i_ u_io_ u_o_*, where *D* is the mean depth and *u_i_*, *u_io_* and *u_o_* are log-normal site, site-by-organ and organ factors with mean 1 (default standard deviations 0.58, 0.17 and 0.27; Supplementary Text, Section 10.3). Site factors above 3 are removed, as by a maximum-depth filter. Optional background artefact sites represent recurrent false positives; they have the same low VAF in every sample, drawn from a gamma distribution (by default with shape 2 and mean 0.023). A site is retained if it is called (in Fig. 6F–I, at least two variant reads) in at least one organ, mimicking the ascertainment of standard somatic callers. Depth tiers are nested filters of one read set, and a site enters a tier only if all organs reach the cutoff.

Fig. 6C–I shows these levels for one set of simulated trees (topology of Fig. 2A, three layers with equal mutation rates, layer contributions (0.10, 0.70, 0.20), unphased bulk samples). Exact transformation shifts layer-fixed variants to 0.05, 0.35 and 0.10 (Fig. 6E). After read sampling at a mean depth of 60, only the peak of L2-fixed variants remains clearly resolvable; L1- and L3-fixed variants are still mostly detected (78% and 93%), but their observed VAFs overlap with the low-frequency variants of all layers and, when present, with background artefacts (Fig. 6F,G). Detection of low-frequency variants is strongly depth dependent. The true VAF at which half of the variant–organ occurrences were detected was about 0.09, 0.03 and 0.013 at mean depths of 20, 60 and 150, respectively (Fig. 6I), so that most private, low-frequency variants generated during organ growth remain unobserved in bulk data. Read-level noise therefore removes information predominantly from the low-frequency and layer-minor parts of the spectrum, which are the parts most informative about organ growth and about the less represented layers.

Together, these observations make simSOMA useful not only for idealized layer-specific phased measurements, but also for designing and interpreting bulk, pseudo-bulk, phased, or unphased sequencing studies in which the observed VAF spectrum is a transformed readout of the same underlying developmental process.

## Conclusion and Discussion

simSOMA provides a modular framework for connecting plant developmental architecture to organ-level somatic VAF spectra. The simulations presented here show that the distribution of somatic variants within and among organs is not determined by mutation rate alone. Instead, VAF spectra emerge from the interaction among ASC niche size, self-renewal turnover, clonal expansion to the SAM periphery, branch-founding bottlenecks, organ precursor number, terminal organ growth, phyllotactic placement, observational sampling, and the topology relating sampled organs.

Several general insights emerge from the simulations. First, parameters controlling ASC lineage persistence have a strong effect on the fixation of somatic variants. Smaller ASC populations and higher turnover increase the probability that mutations become fixed in downstream tissues, whereas larger or more persistent ASC populations maintain lineage diversity for longer. Second, branch founding introduces an additional developmental bottleneck whose strength depends on the number of branch precursors relative to the ASC niche and the sectorized SAM periphery. This makes the effective branch precursor number a key determinant of lineage transmission into lateral branches. Third, organ formation adds a separate source of variation: even when branch-lineage history is fixed, organ precursor number and terminal organ growth determine how inherited and newly arising variants are represented in the sampled organ VAF spectrum. Organ growth produces a neutral 1*/q* tail of private variants whose slope measures the per-division mutation rate in growing organs, which links plant organ spectra to the neutral-tail framework used in cancer genomics and provides a handle on terminal-cell mutation rates. Finally, phyllotactic placement has limited effects on marginal VAF spectra but can leave detectable signals in among-organ sharing patterns when ASC turnover is low.

Although we have focused here on somatic genetic mutations, the same simulation framework can also be applied to somatic epimutations. These are stochastic, mitotically heritable gains and losses of DNA cytosine methylation that arise during cell division and can therefore act as dense lineage markers within developing plant tissues [Johannes and Schmitz, 2019, Chen et al., 2024]. Because genetic mutations and epimutations are transmitted through the same cell lineages, their spatial distribution should be shaped by the same developmental processes. In simSOMA, this means that epimutational accumulation can be simulated by changing the effective mutation rate and target size while keeping the developmental lineage model unchanged. Epimutations occur at substantially higher rates than DNA sequence mutations [Becker et al., 2011, Schmitz et al., 2011, van der Graaf et al., 2015], but over typical developmental time scales recurrent gains or losses at the same site may often be negligible relative to the lineage-level signal [van der Graaf et al., 2015]. An infinite-sites approximation is therefore a useful first implementation, although more detailed versions of the mutation module could incorporate reversible methylation dynamics explicitly where needed.

The parallel to cancer evolution extends beyond the organ tail. In tumours, bulk VAF spectra are routinely interpreted as readouts of clonal expansion, drift and selection [Williams et al., 2016, 2018], and similar reasoning applies to plant organs, with three important differences. First, plant organs are founded by a small, spatially contiguous group of precursors recruited from a structured meristem, so the inherited part of an organ spectrum reflects the sector structure of the meristem rather than a single founding clone. Second, the plant body is a branching hierarchy of such founding events, so variants are shared among organs in proportion to their developmental relatedness, much as subclonal variants are shared among spatially separated biopsies of one tumour. Third, plant cells do not move: lineages are separated by layer and cells remain physically attached to their neighbours, so that precursor recruitment draws on spatially related cells. Somatic evolution in plants is therefore a problem in the population genetics of spatially structured cell populations, whereas many models of tumour evolution assume well-mixed populations. Methods developed for multi-region tumour sequencing [Yu et al., 2020] therefore cannot be transferred directly; they would need to account for the spatial structure and branching hierarchy that simSOMA represents, and simSOMA can provide the corresponding developmental null model.

The observation-model transformations introduced above extend the use of simSOMA beyond the idealized case in which the simulated developmental VAF is treated as the directly measured quantity. By applying phasing and sampling transformations after the developmental simulation, the same underlying topology and cell-lineage history can be viewed under different empirical measurement scenarios. This separation is useful because it distinguishes two sources of variation that are often conflated in empirical VAF data: variation generated by development, and variation introduced by how tissue is sampled and how alleles are counted. It also means that the developmental effects studied here can be re-examined under different observational constraints without rerunning or redefining the core developmental model.

The read-level model makes this separation quantitative. At depths typical of plant somatic-mutation studies, much of the low-frequency and layer-minor signal is lost, so the choice of sequencing depth, layer enrichment and number of sampled organs determines which developmental parameters can be resolved. simSOMA can therefore be used to evaluate sequencing designs before data are generated.

The explicit level of description of simSOMA also has limitations. First, a weak bottleneck requires few amplification divisions between niches (small *C/m*), so that bottleneck strength and the mutational input during amplification are coupled. Second, branch and organ founding currently draw on the same amplified periphery. Precursors that divide less than the surrounding organ-forming cells, as reported for the detached-meristem pathway [Burian et al., 2016], therefore cannot be represented; this would require event-specific amplification. Third, simSOMA simulates SAM layers as independent lineages that share the plant topology and the positions of developmental events, but not cells. These spectra therefore do not capture processes such as cell-lineage exchange between histological layers, which occurs occasionally in angiosperms and more frequently in gymnosperms [Esau, 1965], because the generative model does not include explicit lineage transitions between layers. Studying layer exchange would require a future multilayer extension in which transitions between layers are part of the developmental simulation itself. Such an extension would also require explicit assumptions about where and when layer exchange occurs, for example during axial meristem formation, during pre-branching clonal expansion, or during re-establishment of a new ASC population in a lateral shoot. Because little is known empirically about the timing and frequency of such exchanges, these assumptions would need to be considered carefully, as exchanges introduced at different developmental stages could have different consequences for the resulting VAF spectra.

The present implementation is deliberately modular. This design is useful because several developmental processes represented in simSOMA remain incompletely resolved empirically. For example, organ formation is currently modeled as balanced clonal expansion from a specified number of organ precursors. More detailed models of organ growth, including spatially explicit cell proliferation or empirically calibrated leaf-development trajectories, could be incorporated as alternative organ modules. Similarly, as improved measurements become available, parameters such as ASC niche size, self-renewal rate, precursor number, organ founder number, or cell-division rate can be fixed or constrained to empirically supported values. This would reduce the dimensionality of simulation studies and sharpen the interpretation of the remaining free parameters.

We have not compared simulated spectra with empirical data in this study, whose aim is to describe the simulator and the developmental and observational processes it isolates. Empirical VAF spectra are typically obtained by whole-genome sequencing of bulk or layer-enriched organ samples, followed by somatic variant calling against a reference genome of the same individual and estimation of each variant’s VAF from its variant and total read counts [Goel et al., 2024, Amundson et al., 2025]; the read-level observation model reproduces this process. A natural extension is to use simSOMA as the forward-simulation engine in a Bayesian or approximate Bayesian computation framework. Previous approaches have shown that somatic mutation data can be used to infer developmental parameters or mutation rates under simplified models [Johannes, 2025b, Tomimoto and Satake, 2026, Grecu et al., 2025, Tomimoto and Satake, 2023]. simSOMA extends this direction by providing a configurable simulator that can generate organ-level allele-count and VAF summaries under richer developmental assumptions. This opens the possibility of fitting developmental parameters directly from sequencing data while accounting for topology, founder sampling, organ growth, and observational transformations. Such fitting will have to take into account that several parameter combinations produce similar bottlenecks: only (*P*_eff_ *−* 1)*m/C* determines the probability of polyclonal founding, and *P_b_* cannot be identified beyond *m*. This highlights a general limit of inferring developmental parameters from genetic data: for branch founding, somatic mutation data constrain only this compound quantity, not its components. Constraining parameters that can be measured by histology or live imaging, such as niche size, would reduce this redundancy.

In summary, simSOMA formalizes the idea that somatic VAF spectra in plants are developmental readouts. By linking rooted plant topology with explicit cell-lineage dynamics, it provides a framework for testing how individual developmental assumptions shape somatic genetic and epigenetic mosaicism. More broadly, the simulator offers a practical tool for building intuition, designing sequencing experiments, and developing future inference approaches for somatic evolution in plants.

## Data and code availability

The simulation code is available at https://github.com/jlab-code/simSOMA, and a tutorial can be found at https://github.com/jlab-code/simSOMA-tutorial. The current release, simSOMA version 0.2.3, is archived on Zenodo (doi: 10.5281/ zenodo.23274014). Figs. 2–5 were produced with the version of the original submission (v0.1.0-beta, commit 4ec934e in the repository); the only later change that affects simulated values, a revised displacement rule, applies to *ρ >* 0 and changed the reported fractions by less than 0.005 in the settings checked (Supplementary Text, Section 4.5). Fig. 6C–I and Supplementary Figs. S1–S4 were produced with version 0.2, whose simulation code is identical to version 0.2.3. Topologies and configuration files for Figs. 2–5 are provided as Supplementary files 1 and 2, and the settings and scripts for Fig. 6C–I and the Supplementary Figures as Supplementary file 3.

## AI Declaration

ChatGPT and Claude were used to assist with software development and testing, to help prepare selected sub-figures, and with text editing. I have reviewed all AI-assisted outputs and take full responsibility for the final content of the manuscript.

## Appendix

**Appendix Table 1:**
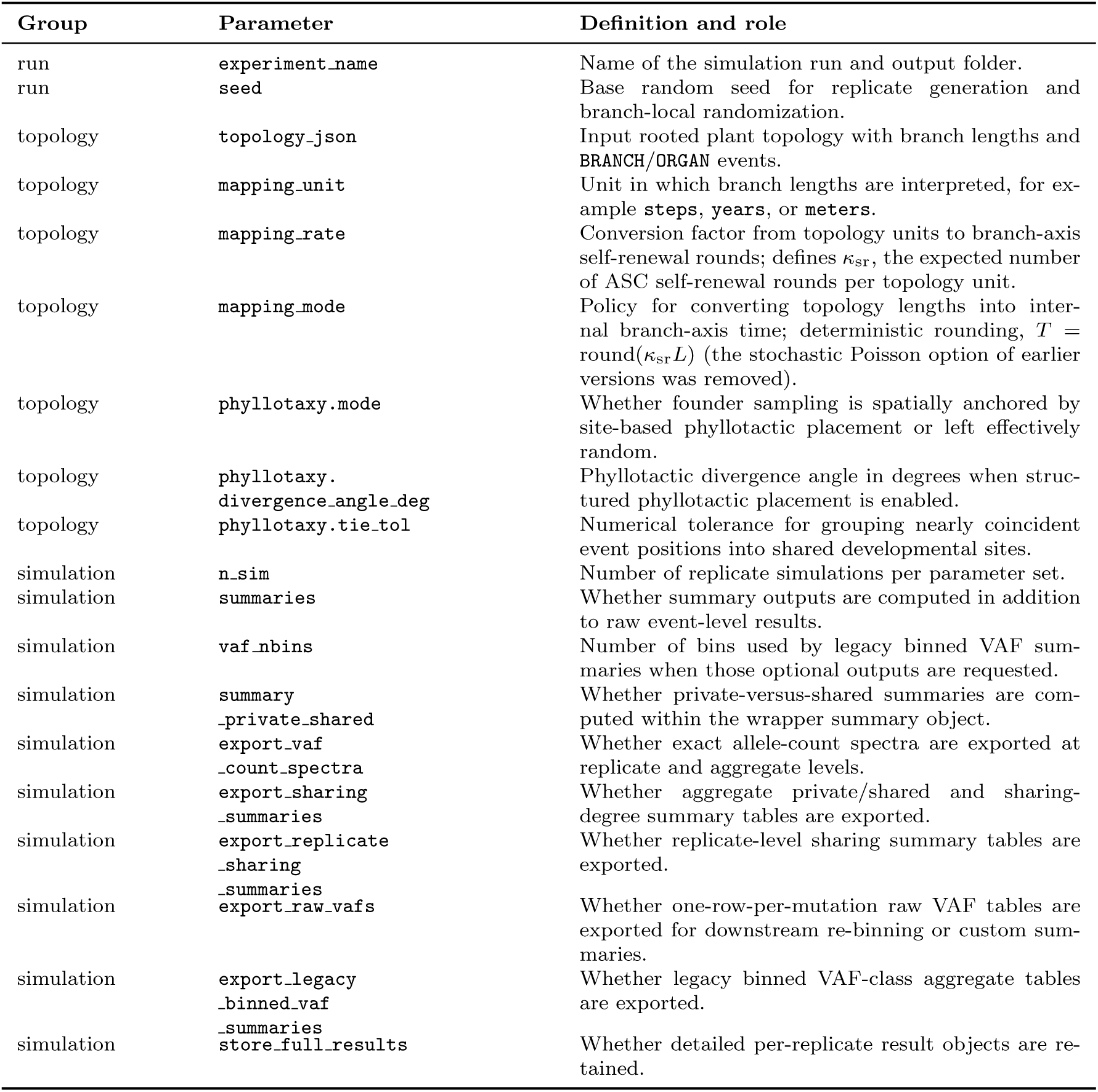
Cross-module and topology-level parameters in simSOMA.

**Appendix Table 2:**
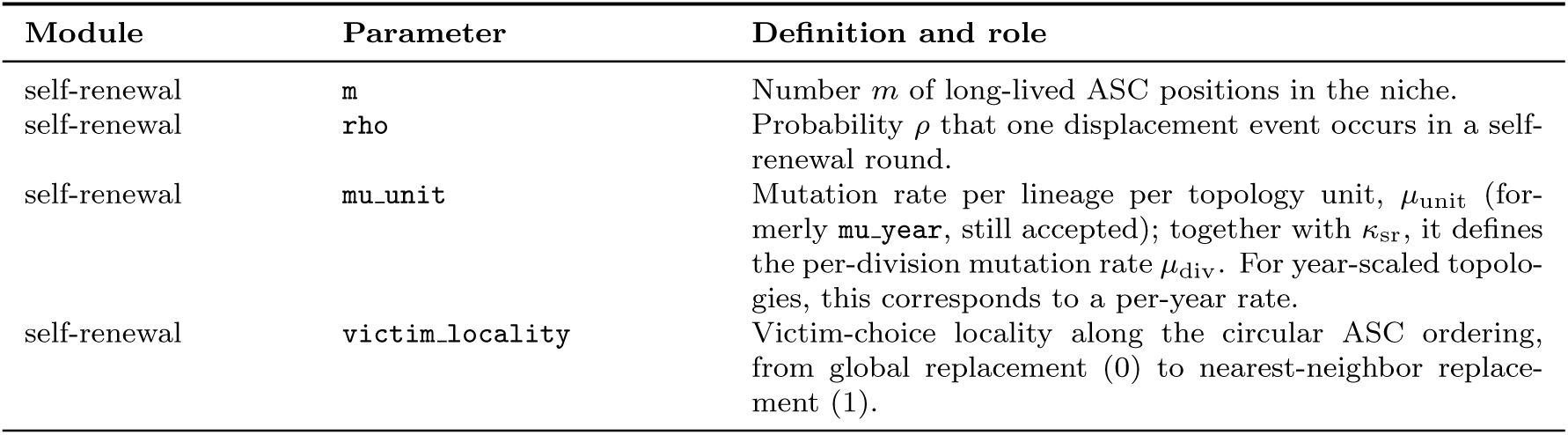
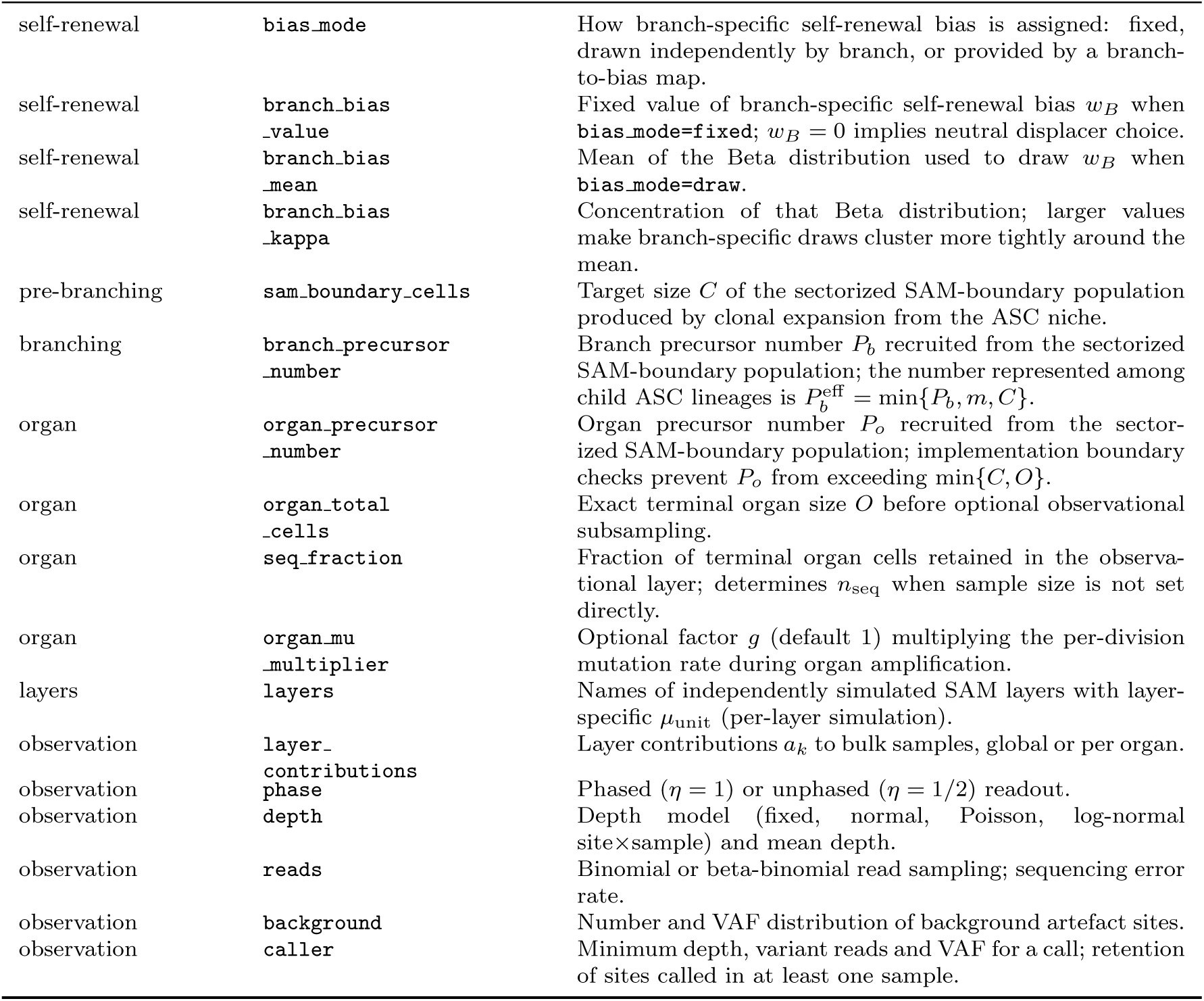
Module-specific parameters in simSOMA.

